# Host population structure and treatment frequency maintain balancing selection on drug resistance

**DOI:** 10.1101/128967

**Authors:** Sarah Cobey, Edward B. Baskerville, Caroline Colijn, William Hanage, Christophe Fraser, Marc Lipsitch

## Abstract

It is a truism that antimicrobial drugs select for resistance, but explaining pathogen- and population-specific variation in patterns of resistance remains an open problem. Like other common commensals, *Streptococcus pneumoniae* has demonstrated persistent coexistence of drug-sensitive and drug-resistant strains. Theoretically, this outcome is unlikely. We modeled the dynamics of competing strains of *S. pneumoniae* to investigate the impact of transmission dynamics and treatment-induced selective pressures on the probability of stable coexistence. We find that the outcome of competition is extremely sensitive to structure in the host population, although coexistence can arise from age-assortative transmission models with age-varying rates of antibiotic use. Moreover, we find that the selective pressure from antibiotics arises not so much from the rate of antibiotic use per se but from the frequency of treatment: frequent antibiotic therapy disproportionately impacts the fitness of sensitive strains. This same phenomenon explains why serotypes with longer durations of carriage tend to be more resistant. These dynamics may apply to other potentially pathogenic, microbial commensals and highlight how population structure, which is often omitted from models, can have a large impact.

---

Despite the enormous burden of antibiotic resistance, the ways in which microbial populations maintain different patterns of resistance are not well understood. The malaria parasite *Plasmodium falciparum* has steadily accumulated resistance to all major drugs used against it [1], whereas *Streptococcus pneumoniae* gains and loses resistance over space and time, at least partly in response to local selection pressure [2,3]. These different dynamics are not yet predictable. It is intuitive that the spread of resistance depends on its genetic determinants and their fitness costs as well as the strength of selection from treatment. But it is also apparent that the spread of resistance changes with the strength of competition between resistant and sensitive strains within hosts, and between resistant and sensitive strains in the host population [4–6]. These interactions can be influenced by treatment practices and host behavior. Effectively managing resistance in microbial populations requires understanding how these dynamics shape the competitive balance between resistant and sensitive strains.

The maintenance of antimicrobial resistance in *S. pneumoniae*, or pneumococcus, presents such a theoretical and practical challenge. Resistant strains seem to be maintained at stable frequencies in different populations, and yet simple models predict that coexistence should be rare, and either sensitive or resistant strains should fix [7]. The patterns suggest balancing selection, whereby coexistence is maintained by negative frequency dependence in the fitness of individual strains. The fraction of strains of *S. pneumoniae* that are non-susceptible to penicillin correlates strongly with local treatment rates among countries in Europe [8–10], provinces in Spain [11], and states in the U.S. [12, 13]. Additionally, periods of high resistance correlate with periods of high local antibiotic usage [14]. These trends suggest that the frequency of non-susceptible strains is driven by treatment, which is consistent with studies showing that amoxicillin therapy disproportionately speeds the clearance of sensitive strains relative to non-susceptible strains [15]. Furthermore, the fact that resistance consistently declines during periods of low antibiotic usage suggests that resistance carries a fitness cost [14, 16, 17]. Sensitive strains tend to outcompete resistant strains in experimentally infected rats in the absence of antibiotics [18]. But the relative advantage of resistant strains in treated hosts, and of sensitive strains in untreated hosts, cannot by itself explain balancing selection for resistance in pneumococcus. Parsimonious transmission models show that resistant and sensitive strains can coexist in only a very narrow range where treatment pressure nearly equals the fitness cost of resistance [7]. Prescription rates of penicillins vary over three-fold between countries in Europe [2, 8], and the high rates of recombination of resistance elements in the pneumococcal genome suggest the fitness cost may vary between strains [19], a hypothesis also consistent with evidence from *in vitro* and animal models [18, 20].

Different forms of competition could mediate selection for resistance and expand the potential for coexistence. In theoretical models, increasing the segregation of sensitive and resistant strains, such as by introducing explicit classes of treated and untreated hosts, slightly increases the range of outcomes in which both strains coexist [7]. Models with greater host population structure similarly predict more coexistence between resistant strains of *Staphylococcus aureus* that are adapted to community versus hospital transmission, although this structure generally does not enable the persistence of sensitive strains [21]. Allowing strains to compete more strongly with themselves than with one another also facilitates coexistence [7]. This kind of competition, mediated by specific immunity, is probably important for maintaining the diversity of pneumococcal serotypes [22]. Serotype variation may be one driver of coexistence, because the fitness benefit of resistance will be larger in serotypes of longer duration of carriage (and thus higher likelihood per carriage episode of antibiotic exposure) [23]. Indeed, prevalence of resistance is higher in more common serotypes, which have longer duration of carriage [24–26]. This mechanism can be generalized: modeling suggests that any phenotype that is simultaneously under balancing selection and associated with variation in duration of carriage will, by epidemiologically generated linkage, be associated with antibiotic resistance [23]. Candidate mechanisms in addition to serotype, such as phage content, have been identified [27], but it is still unclear how much variation in resistance this effect explains, especially within serotypes. Here we focus instead on quantifying the impact of effects related to population structure.

To investigate the forces maintaining balancing selection on resistant and sensitive strains, we modeled the competitive dynamics of pneumococcus from the scale of individuals to communities. Resistant strains of pneumococcus are ubiquitous in human populations and arise repeatedly through recombination, and so we focus on the effects of treatment and host behavior rather than the evolution of resistance de novo. Our results uncover new mechanisms contributing to balancing selection, including an unexpected role of treatment frequency, and explain differences in resistance rates between serotypes. Such insights suggest new areas of focus for managing pneumococcal resistance.

## Results

We used an individual-based model to simulate the transmission dynamics of 25 pneumococcal serotypes in a host population whose age structure and vital rates were similar to those of the United States (Methods). Serotypes differed in their intrinsic fitnesses, with some serotypes having longer duration of carriage and greater within-host competitive ability, and thus higher prevalence, than others. Serotypes occurred as resistant and sensitive strains. Sensitive strains had longer durations of carriage in untreated hosts, and resistant strains had longer durations of carriage in treated hosts, but there was otherwise no difference between strains of a given serotype. Although more than 90 serotypes have been recorded, a smaller number are typically present in any given sample and we have selected 25 as a tractable number that is capable of capturing the observed diversity. Rates of clearance in treated hosts were fixed for both strains based on experimental values [15]. Treatment occurred independently of the host’s carriage status. To mitigate the effects of extinction due to drift in our artificially small population of 100,000 hosts, individuals were exposed to immigrating strains at approximately 0.1% the rate at which they were exposed to strains from within the population. We examined the consequences of varying the fitness cost of resistance, defined as the percent reduction in resistant strains’ duration relative to sensitive strains’ in untreated hosts, and the intensity of treatment, which was normalized to the U.S. rate of penicillin use. For the different models, we examined the range of fitness costs in which resistant and sensitive strains could coexist.

### High antibiotic pressure alone can only select for resistance with modest cost

We started with a randomly mixing population in which there is no difference among hosts in the probability of receiving treatment. Hosts vary in their treatment statuses over time, creating a shifting pool of habitats favorable to the survival and transmission of either sensitive or resistant strains. In simpler models, including discrete treated and untreated host compartments has slightly promoted coexistence [7]. In our simulations, increasing the rate of antibiotic treatment predictably increased the fraction of resistant pneumococci. As expected, these increases shrank as the cost of resistance grew (Fig. 1). Over the range of treatment rates observed in Europe (corresponding to treatment levels of roughly 0.5-1.5 in our model), only very small fitness costs (≤ 2%) were compatible with the coexistence of resistant and sensitive strains. This assumes a broad definition of coexistence, where resistant and sensitive strains each have frequencies between 2% and 98%. We further require for coexistence that the fraction of resistant strains increases at least 10% over this range of treatment rates (e.g., if 40% of strains are resistant at the 0.5 treatment level, at least 44% should be resistant at the 1.5 treatment level) to match the observed positive correlation between resistance and treatment [8–13]. Because epidemiological and experimental evidence suggests the cost may exceed 2% [14, 18], we conclude that a randomly mixed, equally treated population cannot maintain stably coexisting penicillin-sensitive and resistant pneumococci given mean penicillin prescription rates, despite the natural discretization of treated and untreated individual hosts.

**Figure 1:**
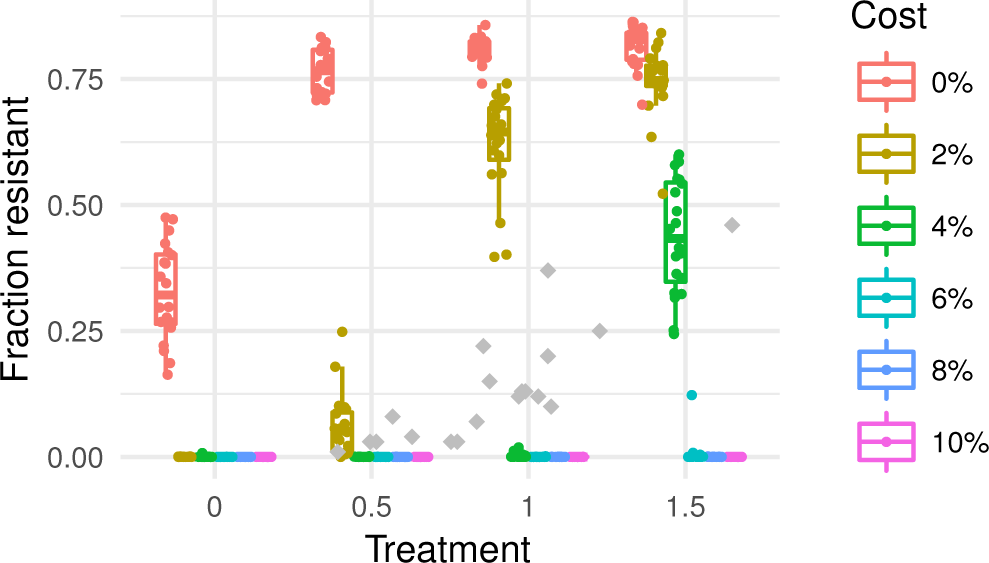
Fraction of all pneumococci resistant by treatment level and fitness cost for the model with identical treatment rates by age and random mixing. Each colored point shows the mean from one replicate simulation. Gray points show resistant fractions observed in European countries [2].

### Structure from transmission and treatment promote coexistence

We next evaluated the impact of three additional forms of host population structure, all biologically supported, that might promote the coexistence of resistant and sensitive strains at greater fitness costs. People preferentially contact others in their age group, and so we simulated transmission dynamics with age-assortative mixing drawn directly from contact surveys [28]. Assortative mixing slightly increases the rate of coexistence when resistance carries a cost (Fig. 2).

**Figure 2:**
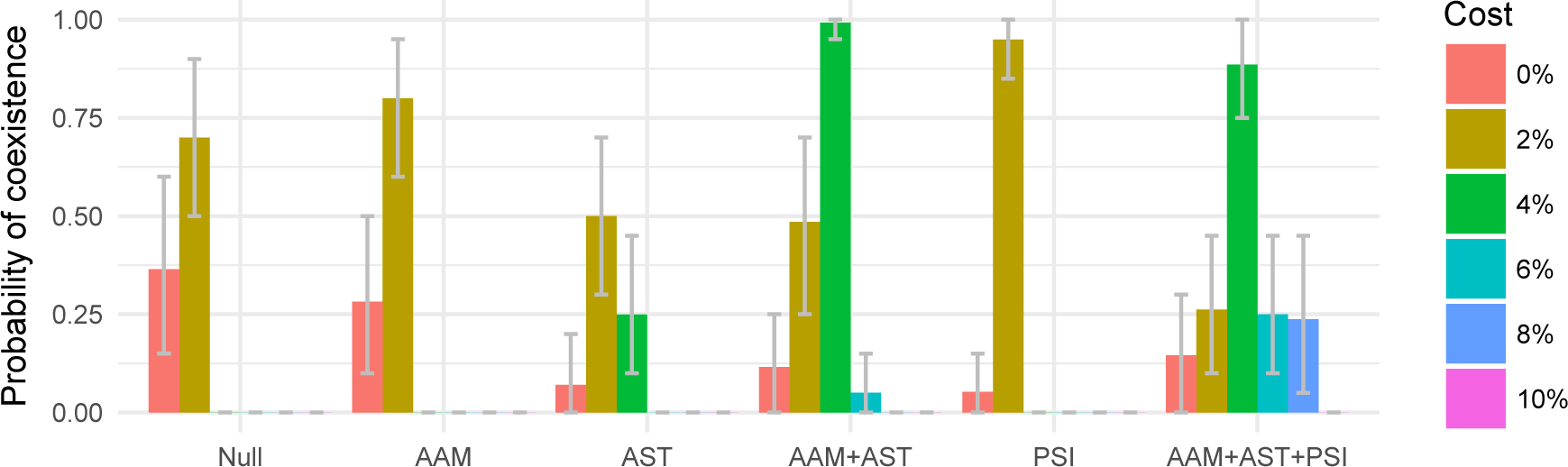
Fraction of replicate simulations producing coexistence under different models and fitness costs. Parameters are identical between replicates. Coexistence is defined as resistant fraction between 2% and 98% and a 10% increase in this fraction as the treatment intensity varies from 50% (treatment=0.5) to 150% (treatment=1.5) of typical rates. One thousand sets of randomly selected simulations at the three treatment levels were examined (e.g., the data associated with treatment = 0.5, 1.0, and 1.5 in Fig. 1). Error bars show 95% of the distribution of coexistence means from bootstrapping 1000 times over 20 triplets of runs at different treatment rates. AAM = age-assortative mixing, AST = age-specific treatment, PSI = pseudo-spatial immigration.

Antibiotic prescription rates are higher in children than adults, especially for respiratory tract infections. Keeping the total rate of antibiotic usage in the population constant, we varied the treatment rates by age group to match the distribution of penicillin prescriptions in U.S. populations (Fig. S1). Without treatment, the model predicts higher carriage rates in children than adults due to their differences in immunity [22]. Shifting treatments to young children predictably increases the fitness of resistant strains, increasing the overall fraction of resistant pneumococci in the population as a whole and also differences between the frequency of resistance in children and adults (Fig. 3). Carriage rates remain higher in children than adults even though children receive proportionately more antibiotics (Fig. S2). Compared to both the null model and a model of age-assortative mixing alone, a model with age-specific treatments extended the range of fitness costs compatible with coexistence of resistant and sensitive strains (≤ 4%; Fig. 2).

**Figure 3:**
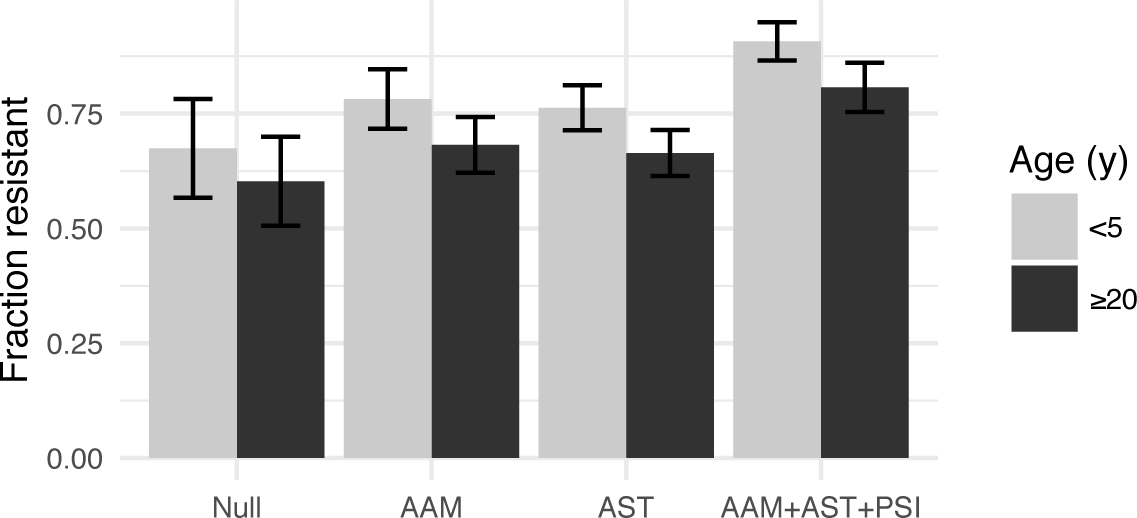
Fraction of pneumococci resistant in young children (*<* 5 y) and adults (≥ 20 y), for each model, with treatment at U.S. rates and a 2% cost. Error bars give the SD of the means of the replicates. AAM = age-assortative mixing, AST = age-specific treatment, PSI = pseudo-spatial immigration.

We next considered the impact of host population structure over a larger scale. In the simulated population of 100,000 hosts, the less common serotypes occasionally go extinct. By default, we assume that all immigrating strains have a 1% probability of being resistant, regardless of serotype. A more plausible assumption is that immigrating strains have the same probability of being resistant as a random strain of that serotype in a population with typical treatment rates (Methods). With “pseudo-spatial immigration,” the range of costs supporting coexistence is narrower than with age-specific treatments, although coexistence is more likely at a 2% cost (Fig. 2). Because immigrating strains are involved in approximately 0.1% of colonization events on average, immigration’s main effect on coexistence is to reseed strain populations that have gone extinct. Small changes in the probability of resistance among immigrants can nonetheless exert a slight effect on the probability of coexistence at different costs.

Combined, these forms of host population structure increase the range of fitness costs that allow coexistence and the frequency of coexistence for any cost (Fig. 2). Increased segregation of treated and untreated hosts through age-assortative mixing and age-specific treatment expands the range of tolerated costs. Pseudo-spatial immigration increases the frequency of coexistence across a wider range of costs. These results were robust to some uncertainty in the duration of carriage of resistant strains in treated hosts (Figs. S3, S4).

### Co-transmission has little effect on coexistence

Contacts with multiply colonized hosts create correlations in the risks of contact (and thus colonization) with different strains. Correlated transmission of groups of resistant or sensitive strains might promote their persistence in treated and untreated hosts. In the default model, by contrast, the risk of contacting a strain is identical for all hosts, or for all hosts of the same age with age-assortative mixing. We investigated the impact of correlated risks by simulating contacts between individual hosts, allowing the colonized host to “challenge” the recipient host with each of the colonized host’s strains (Methods). Models with co-transmission modestly affected the frequency of coexistence (Fig. S4). Compared to the null model, models with cotransmission did not expand the range of costs supporting coexistence. However, when there was already host population structure, cotransmission could slightly increase the frequency of coexistence at the same range of costs if multiply colonized hosts were assumed to be equally infectious as singly colonized hosts (that is, if the probability of contacting each of the strains was divided by the total number of strains carried by the donor host). The minor impact of cotransmission is unsurprising because over 80% of hosts in these models are colonized with only one strain.

### Common serotypes are more resistant

When resistant and sensitive strains coexist, resistance is more common in the most fit serotypes (Fig. 4), with the most fit ones sometimes showing fixation of resistance and the least fit showing fixation of sensitivity. This pattern changes little with age (Fig. S5). Serotypes in our model all have the same transmission rate, but they vary in two fitness components: their maximum expected durations of carriage and their ability to exclude other colonizing strains, and thus in prevalence. The most fit serotype persists, on average, 220 days in naive, untreated hosts, whereas the rarest serotypes persist only 25 days. More fit serotypes thus are more likely to experience treatment during a carriage episode, and thus experience greater selection for resistance [23]. This pattern matches trends in pneumococcal epidemiology [24–26, 29]. These results show that not only does duration modulate whether sensitive or resistant strains are most fit for a given serotype, as previously shown using a simpler model [23], but in the presence of other mechanisms promoting coexistence, a range of durations can produce a range of prevalence of resistance within serotypes, from zero for low-fitness types, through coexistence to 100% for the highest-fitness ones. Resistance is higher in children than adults (Fig. 3) because children tend to be infected by the more resistant serotypes due to their longer duration of carriage (specifically in children) and to the fact that most adults have experienced these common serotypes and are partially immune to acquiring them; both effects are amplified by age-assortative mixing such that children’s exposures are disproportionately to other children (Fig. S6). Selection for resistance in these serotypes overall also increases slightly with age-assortative mixing and age-specific treatments.

**Figure 4:**
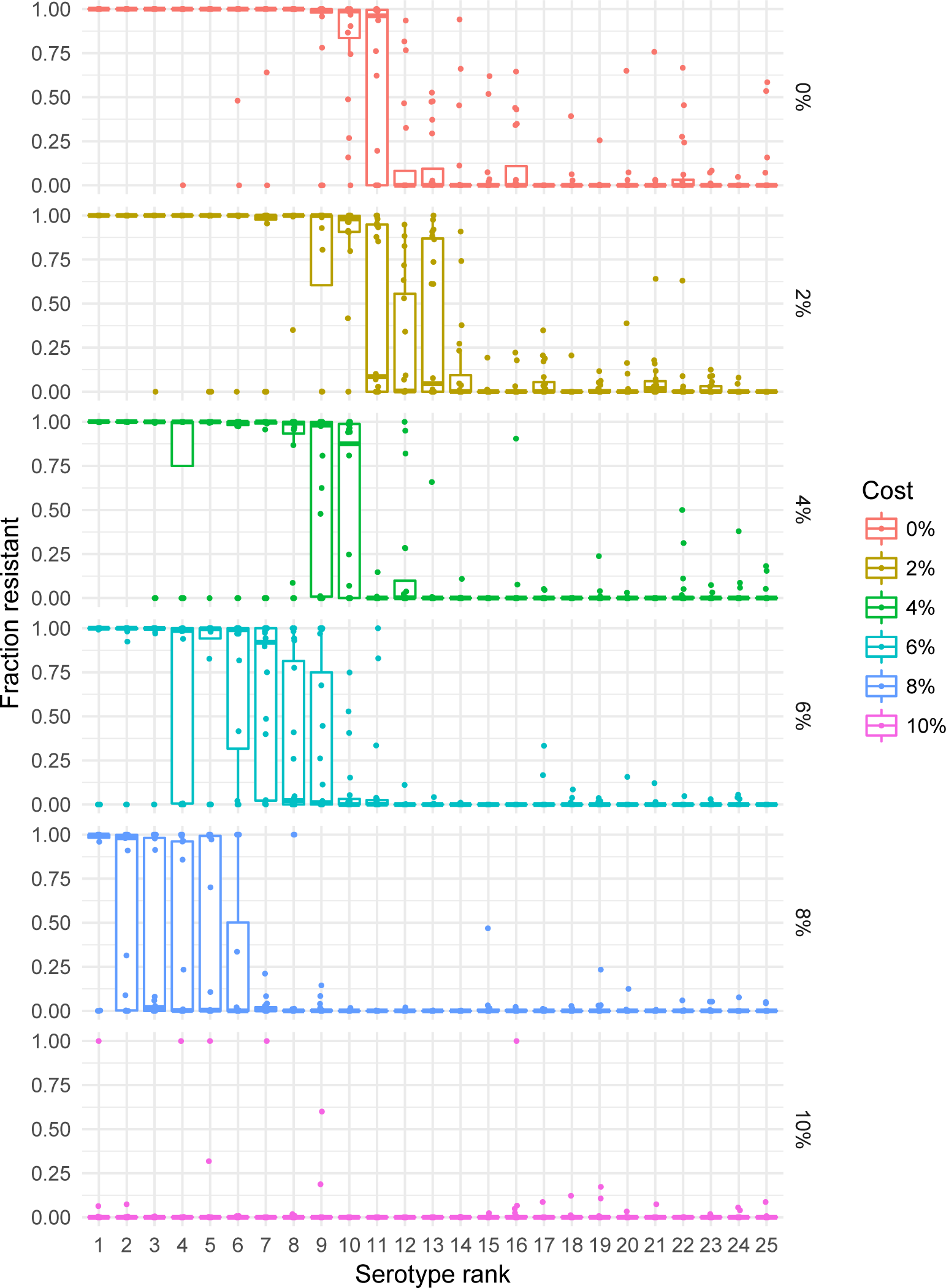
Resistance in young children (*<* 5 y) by serotypes’ fitness ranks for different costs of resistance in the model with age-assortative mixing, age-specific treatment, and pseudo-spatial immigration. Treatment is held at 1.0, equivalent to U.S. rates. Each point shows the mean fraction of each serotype that is resistant from one replicate simulation. Means were obtained by averaging the last 50 years.

### Treatment incidence more important than prevalence for promoting resistance

For the same reason that treatment disproportionately affects the fitness of serotypes with long duration, increasing the incidence of treatment episodes—but leaving total antibiotic use for each age unchanged—could promote resistance in the pneumococcal population as a whole: shorter intervals between treatment events increase the number of colonizations with sensitive strains that can be disrupted, and thereby increase selection for resistance across all serotypes. Instead of a 10-day treatment duration, we simulated 2.5- and 5-day treatment durations while quadrupling and doubling the expected number of treatment events per capita, respectively, and also a 20-day treatment duration while halving the treatment frequency. Total antibiotic use still varied by age as before (Fig. S1), but for each age, the mean duration determined whether there were more short treatments or fewer long treatments. Increasing the frequency of treatment reduced the mean fitness of sensitive strains, allowing resistant strains to persist at a high fitness costs, even when antibiotic use was low. It also increased the fraction of replicate simulations producing coexistence across all fitness costs (Figs. 5, S4).

**Figure 5:**
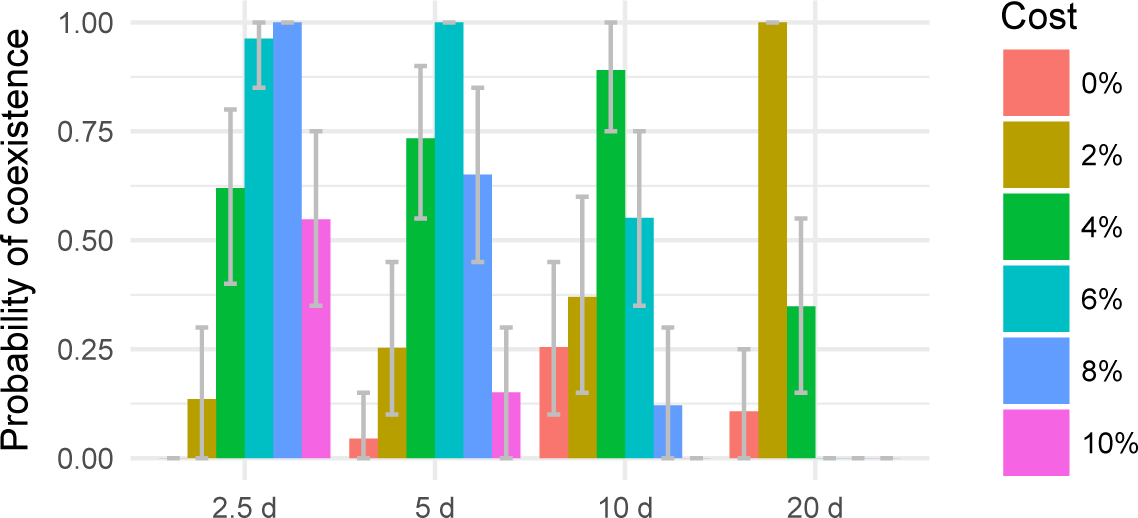
Fraction of replicate simulations producing coexistence for different treatment durations, which correlate inversely with treatment frequency. Parameters are identical between replicates. The mean prevalence of treated hosts is held constant across models, so 5-day treatment durations are associated with twice the number of treatment events as the 10-day default, and 20-day treatment durations are associated with half as many. All models include age-assortative mixing, age-specific treatments, pseudo-spatial immigration, and cotransmission with equally infectious hosts. Coexistence is defined as resistant fraction between 2% and 98% and a 10% increase in this fraction as the treatment intensity varies from 50% to 150% of typical rates. Error bars show 95% of the distribution of coexistence means from bootstrapping 1000 times over 20 triplets of runs at different treatment rates.

## Discussion

It is not easy to justify balancing selection for resistant and sensitive strains of pneumococcus from first principles. Simple models predict that sensitive strains should generally exclude resistant strains when fitness costs of resistance are high, and resistant strains should exclude sensitive strains when the cost is low [7]. This explanation is unsatisfying given the observed range of frequencies at which both resistant and sensitive strains appear to coexist, and given the likely variation in resistance cost. We find that host population structure is critical to strain persistence, as it creates semi-protected habitats (niches) in which each strain has a fitness advantage. Our analysis highlights another aspect that is missing from some simple models. Strain fitnesses depend not only on the overall prevalence of antibiotic treatment in the population but also on the frequency of treatment episodes. This dependence helps to explain why common serotypes, which have long carriage durations, tend to be resistant. An aspect not considered in this model is that shorter courses may be given with higher doses [15], which can shift selection against susceptible and intermediately resistant strains, and in favor of more highly resistant strains [30]. This model also considers resistance a binary, rather than continuous, trait. Nonetheless, our finding suggests that all else being equal, resistance evolution should be more sensitive to the number of treatment courses than to the number of treatment-days of antimicrobial use. Overall, our results suggest that managing resistance in pneumococcus should include consideration of treatment frequency, not just its prevalence, as well as the dynamics of transmission and treatment in core groups such as children.

Aggressive chemotherapy can prevent the de novo emergence of resistance strains if drug-sensitive strains are suppressed sufficiently rapidly [4, 5], but when relatively fit resistant strains are already in circulation, aggressive chemotherapy can remove drug-sensitive competitors and speed the selection and emergence of resistant strains. Selection in our model proceeds by the latter mechanism. By rapidly clearing sensitive strains, treatment reduces their fitness advantage in transmission, promoting the growth of resistant strains in the community. This is consistent with epidemiological studies, which show that the clearance of sensitive strains following treatment appears to have a stronger effect on individual risk of carriage than competitive release of resistant strains [31]. In other words, although treated hosts are at increased risk of colonization with resistant types, the fraction of resistant strains is mostly driven by changes in the abundance of sensitive strains. A corollary is that the prevalence of treatment should correlate negatively with the overall prevalence of pneumococcus in the community. This pattern appeared in the model (Fig. S7) but has not, to our knowledge, been investigated in natural settings.

Resistant fractions above 60% are rarely observed in large populations [2, 8], suggesting that our models cannot easily explain observed patterns: our simulated pneumococci are consistently too strongly affected by a threefold variation in selective pressure (Figs. S3, S8). There are several additional mechanisms, not included in our model, that may produce coexistence over a wider range of parameter values in reality than our model can capture. With age-assortative mixing and age-specific treatment, the spread of resistant strains is mostly limited by the fitness cost of resistance and the intensity of treatment. Relative to natural populations, our simulated hosts have little spatial and demographic structure. Households, schools, and regional transmission all influence pneumococcal dynamics. Because these structures amplify the segregation of treated, high-transmitting hosts (especially children), they should facilitate constrained spread of resistant strains. This is especially likely if antibiotic treatment tends to follow infections by respiratory viruses, which have spatially localized epidemics. Spatiotemporal variation with migration might explain why resistance rarely fixes in the most common serotypes, as our model sometimes predicts. A separate potential mechanism promoting coexistence is genetic linkage between resistance to *β*-lactams and other antibiotics [32]. Use of other antibiotics would thus select inadvertently for resistance to *β*-lactams. This may be an important factor driving resistance: trimethoprim-sulfamethoxazole and cephalosporin use can be better predictors of penicillin non-susceptibility in individual hosts and communities than penicillin use [11, 12, 33]. The effective rate of antibiotic treatment may thus be higher than predicted from penicillin prescriptions alone (which comprise approximately one-third to two-thirds of total antibiotic prescriptions in European ambulatory settings [10]), promoting the spread of resistant strains and increasing the range of tolerable fitness costs. Finally, if resistant elements induce adaptive immunity, or if they are linked with structures that do, strong intrastrain competition could promote coexistence [7]. In the same vein, variation in the capsular locus and other loci that influence the duration of carriage, such as phage [27], and that are also under negative frequency dependent selection is expected to promote coexistence through epidemiologically generated linkage [23]. Carriage duration determines the benefit of resistance, and selection for different durations indirectly selects for varying degrees of resistance.

The purpose of this work has been to explore the effect of known heterogeneities in population structure, which we show may explain some of the observed coexistence; our approach focused on isolating individual model components to understand each component in isolation, and in combination. Combining the effect of population structure with other mechanisms, such as genetic variation in carriage duration beyond the capsular locus [23] and genetic linkage between resistance to different antibiotics, may fully explain coexistence, but will also likely result in complex models that are hard to understand or validate. A parsimonious approach will focus on combining validated and quantified effects with known impact on antibiotic resistance prevalence.

This work demonstrates the importance of seemingly fine details of model structure in developing expectations about competing strains. Previous models based on ordinary differential equations considered several elements of our model separately, but they omitted key features. We suspect that the most important are the discretization of treatment to individual hosts (similar to a treated compartment) and individual periods, as well as the fine population and age-structure from age-assortative mixing and age-specific treatment rates. We show the importance of treatment frequency, not merely treatment rate. The simultaneous colonizations with multiple strains allowed by our model affect competition within hosts; simultaneous transmission of dual infections was previously shown to be important for coexistence [7]. Investigating the dynamical differences between seemingly parsimonious models, and identifying the rates that determine, e.g., the strength of competition in different scales, will be important for rationally managing resistance.

## Methods

The first section contains a general overview of the model, whose core has been described previously [22]. Later sections describe components in depth. The parameters are in Table 1.

**Table 1:** Parameters in the models.

| Parameter | Description | Value | Rationale |
| --- | --- | --- | --- |
| $\beta$ | transmission rate | $0.045 \text{ d}^{-1}$ | see text |
| $w$ | per capita immigration rate per serotype | $1.25 \times 10^{-7} \text{ d}$ | see text |
| $p_z$ | probability immigrating strain of serotype $z$ is resistant | 1% | see text |
| $\alpha$ | age-assortative mixing weights | ... | [28] |
| $\sigma$ | susceptibility reduction from serotype-specific immunity | 0.3 | [22, 36] |
| $\gamma(z)$ | intrinsic duration of sensitive strain carriage in untreated host | 25 to 220 d | [22, 37, 38] |
| $\kappa$ | minimum duration of carriage in untreated host | 25 d | [22, 38] |
| $\epsilon$ | shape parameter for non-specific immunity | 0.25 | [22, 37] |
| $\xi$ | fitness cost of resistance | [0.9, 1] | varied |
| - | sensitive strain clearance rate with treatment | $1/4 \text{ d}^{-1}$ | [15] |
| $\gamma_r$ | ratio of resistant to sensitive strains' durations with treatment | 4 | varied |
| $\mu_{\max}$ | reduction in susceptibility from carrying fittest serotype | 0.25 | [22, 39] |
| - | per capita treatment rate (unnormalized) | $0.354 \text{ y}^{-1}$ | varied |
| - | mean treatment duration | 10 d | [15] |
| - | SD of treatment duration | 3 d | ... |
| - | minimum interval between treatments | 1.5 d | ... |
| $N$ | number of hosts | 100,000 | ... |
| - | PMF of age at death | ... | [22, 40] |
| - | initial fraction of hosts immune | 0.5 | see text |
| - | initial fraction of hosts colonized with each serotype | 0.02 | see text |
| - | initial fraction of colonizing strains resistant | 0.5 | see text |

### Overview

We used an individual-based model that tracked the carriage histories and ages of hosts over multiple generations [22]. Twenty-five serotypes of pneumococcus, each with two resistance phenotypes (resistant or sensitive), were simulated. All serotypes had the same transmission rate, but they differed in their durations of carriage and ability to exclude other serotypes when colonizing a host. These two traits were positively correlated, so there was a consistent ranking of serotypes’ fitnesses. Hosts gained serotype-specific and non-serotype-specific immunity, the former reducing susceptibility to future colonizations and the latter incrementally shortening the duration of carriage. Immunity thus depended on the total number of prior pneumococcal colonizations and their serotype composition, and it was independent of strains’ antibiotic sensitivities.

Antibiotic resistance could have a cost. For each serotype, resistant strains had a duration of carriage in untreated hosts that was some constant fraction *ξ* of the sensitive strains’ durations (in untreated hosts) [18]. Values of *ξ <* 1 imply an intrinsic cost of resistance. Because the cost of resistance is uncertain, a range of values of *ξ* was tested, and the cost was expressed as a percentage ((1 *− ξ*) *×* 100%).

Both resistant and sensitive strains were negatively affected by antibiotic treatment, but sensitive strains were impacted more. Clearance rates in treated hosts have been estimated as 0.32 day^−1^ for sensitive strains and 0.14 day^−1^ for resistant strains [15]. We fixed the mean clearance rate of sensitive strains in treated hosts 0.25 day^−1^ (a mean duration of 4 days) and assumed resistant strains lasted four times as long. This clearance rate of resistant strains in treated hosts is lower than empirical estimates, but if resistant strains were cleared much faster, antibiotic use dramatically depressed carriage prevalence in the population. Such a trend has not been reported.

To avoid spurious results from stochastic extinctions in our finite host population, we allowed each serotype to have a very small rate of immigration *w*. The rate of immigration *w* was set equivalent to one individual in the population being exposed every 80 days to another individual outside the population carrying each serotype-strain combination. By default, immigrating strains had a 1% probability of being resistant, *p_z_* = 0.01, regardless of serotype *z*. With pseudo-spatial immigration, ten populations with average antibiotic usage rates (treatment = 1.0) were simulated for 150 y with *p_z_* set individually for each serotype to the current fraction of colonizations resistant in the population, or *p_z_* = 0.01 if no colonizations for that serotype were present. The fraction of resistant strains in each serotype was then averaged over the last 50 years and over all simulations to calculate the long-term fraction of resistance in each serotype. These serotype-specific values of *p_z_* were then used in simulations with other rates of antibiotic use.

### Transmission

At discrete time steps, the force of colonization on a host for each serotype *z* and strain *i* (i.e., a resistant or sensitive strain) in the simplest model was computed following

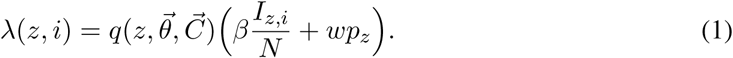

The term 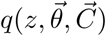 is the host’s susceptibility, or probability of acquiring serotype z, contingent on the vectors of past colonizations 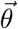 and current carriage 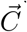. Vectors 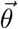 and 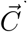 are both indexed by serotype. Entries of 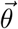 count the number of times the host has cleared each serotype, and 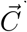 counts the number of current colonizations with each serotype. The contact rate, *β*, is shared by all serotypes. The effective fraction of hosts colonized with serotype *z* and strain 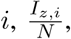 equals the number of colonizations with (*z*, *i*) in the population divided by the population size *N*. This representation means that hosts who are colonized with a single strain of serotype *z* are counted once, and hosts harboring more than one strain of serotype *z* are counted multiple times; this formulation avoids biasing the model toward coexistence [34].

In the model that includes age-assortative mixing, the force of colonization with serotype *z*, strain *i*, on a host of age *a* is computed by

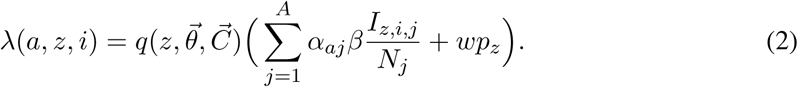

Here, *α_aj_* weights contacts by individuals of age *j* to individuals of age *a*. The matrix *α* of those weights is obtained from normalizing data from physical contact surveys from the United Kingdom and has single-year resolution [22, 28].

### Co-transmission

Two alternative modes of transmission test the impact of correlations in colonization risk with different strains. In each time step, the expected number of potential transmission events in the population is calculated as *βI*, where *I* gives the total number of people with any pneumococcal colonization. For each contact, a random colonized host and random other host are selected. Under the “equally infectious hosts” assumption, the receiving host is challenged with each colonization in the colonized (donor) host with probability 1*/n*, where *n* is the number of colonizations. Under the “equally infectious colonizations” assumption, the challenge probability for each colonization is 1, i.e., the rate of transmission is unaffected by the total number of colonizations in the donor. The overall transmission rate is effectively higher in the second model. The probability of colonization given challenge is determined by the receiving host’s susceptibility.

### Host susceptibility to colonization

The host’s susceptibility to a serotype is reduced if the host is currently colonized with pneumococcus and further reduced if the host has previously carried (i.e., cleared) that serotype. Assuming *τ*(*z*) = 0 if *θ_z_* = 0 (i.e., the host has not previously cleared serotype *z*) and *τ*(*z*) = 1 otherwise, the susceptibility 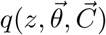 is given by

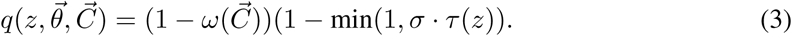

where 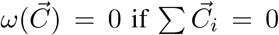 (i.e., if the host is not carrying pneumococcus). 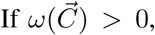 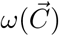 gives the reduction in susceptibility to an invading serotype caused by immediate exclusion by the most fit resident. If 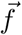 is the vector of fitness ranks of the carried serotypes (ranked against all serotypes), with min 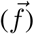 denoting the most fit carried serotype, the reduction in susceptibility is given by

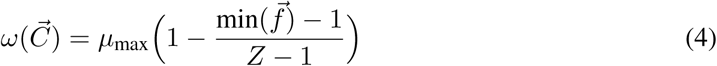

where *Z* gives the total number of serotypes. Equation 4 scales the resistance to acquisition of a new serotype, such that the most fit serotype reduces acquisition by a fraction *µ*_max_, and this resistance declines to zero for the least fit serotype. This value has been estimated to be approximately 0.3 [22]. With cotransmission, strains acquired during the same contact event do not affect 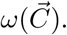

Serotype-specific immunity is denoted by *σ*; it gives the fraction reduction in susceptibility to colonization based on any previous exposure to that serotype.

### Duration of carriage

When a host becomes colonized with a strain of serotype *z*, a duration of carriage for that colonization event is drawn from an exponential distribution with mean *ν_s_*(*z*). The mean depends on the serotype-specific duration of carriage in the absence of immunity, *γ*(*z*), the host’s treatment status, and the total number of past colonization events, independent of serotype. In an untreated host, the duration of carriage of a sensitive strain is calculated as

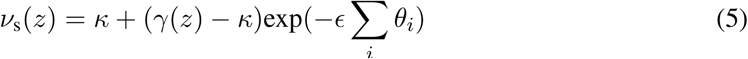

The minimum duration of carriage, which is constant across serotypes, is given by *κ*. Parameter *∊* sets the rate at which non-specific immunity accumulates and was previously fitted [22]. In untreated hosts, resistant strains face a fitness disadvantage, and their carriage duration *ν*_r_(*z*) = *ξ*_s_(*z*).

In a treated host, both sensitive and resistant strains have constant daily rates of clearance that are independent of their serotype. Sensitive strains clear, on average, every 4 days [15], and resistant strains clear every 16 days.

### Treatment

Hosts were randomly assigned treatments at birth in accordance with the age-specific rates of treatment. Thus, treatments did not depend on carriage status. The duration of treatment was drawn from a normal distribution with a mean of 10 days and standard deviation of three days. Treatments were required to be spaced at least one day apart.

Age-specific treatment rates were calculated by multiplying the annual number of antibiotic prescriptions for each age class in the U.S. in 2005-2006 by the fraction of antibiotic prescriptions from acute respiratory tract infections that were penicillins [35]. This yielded an age-specific estimate of the annual rate of age-specific penicillin prescriptions. Multiplying by the standard duration of treatment (10 days [15]) yielded an estimate of the daily per capita prevalence of penicillin use. This estimate was lower than measurements obtained from measures based on antibiotic sales [8] and was scaled to match. This U.S. rate became the reference standard to which other treatment rates were normalized (Fig. S1).

### Demography

At birth, a host’s lifespan was drawn stochastically from the probability mass function of host lifespans in the U.S. Each death caused a birth, so that the population size did not change.

### Simulations

Epidemiological dynamics were simulated for 150 years, and the force of colonization was calculated daily. Simulations started with approximately 2% of hosts colonized with each serotype, 50% of hosts with serotype-specific immunity, and 50% of colonizations resistant. The abundances of resistant and sensitive strains were averaged from annual samples from each of the last 50 years of simulation. For each parameter combination, twenty replicate simulations were performed.

## Authors’ contributions

Conceived the study: SC, CC, WH, CF, and ML. Designed the study: SC, CF, and ML. Developed the software: SC and EB. Performed the simulations and statistical analyses: EB. Created the figures: EB. Wrote the manuscript: SC. All authors edited the manuscript and gave final approval for publication.

## Data accessibility

Code for the simulations can be freely downloaded at https://github.com/cobeylab/pneumo-resistance.

## Acknowledgements

This work was completed in part with resources provided by the University of Chicago Research Computing Center.

## Supporting Information

**Figure S1:**
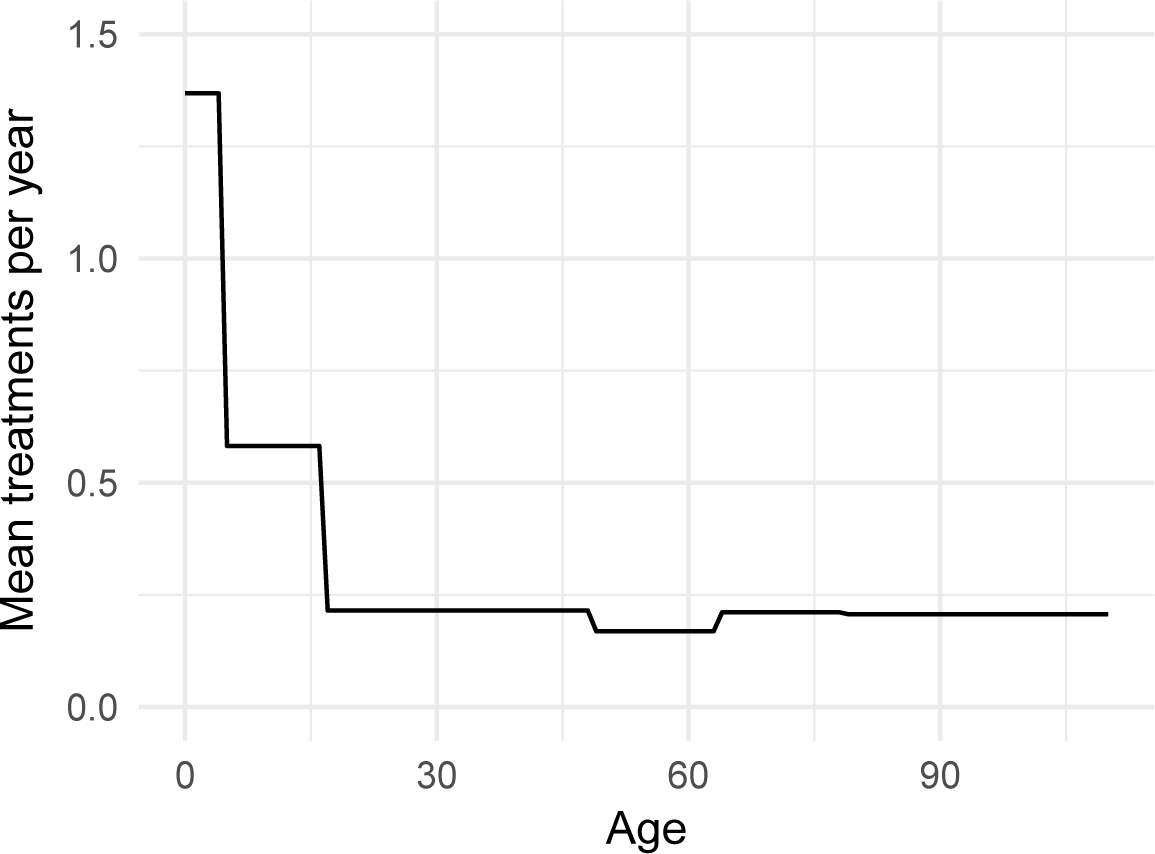
Mean number of treatments per year for each age group. The default treatment duration was 10 days (Table 1). When the treatment duration was varied, the total antibiotic use (number of treatment days *×* number of treatments per year) was kept constant for each age group, and the number of treatments per year was rescaled.

**Figure S2:**
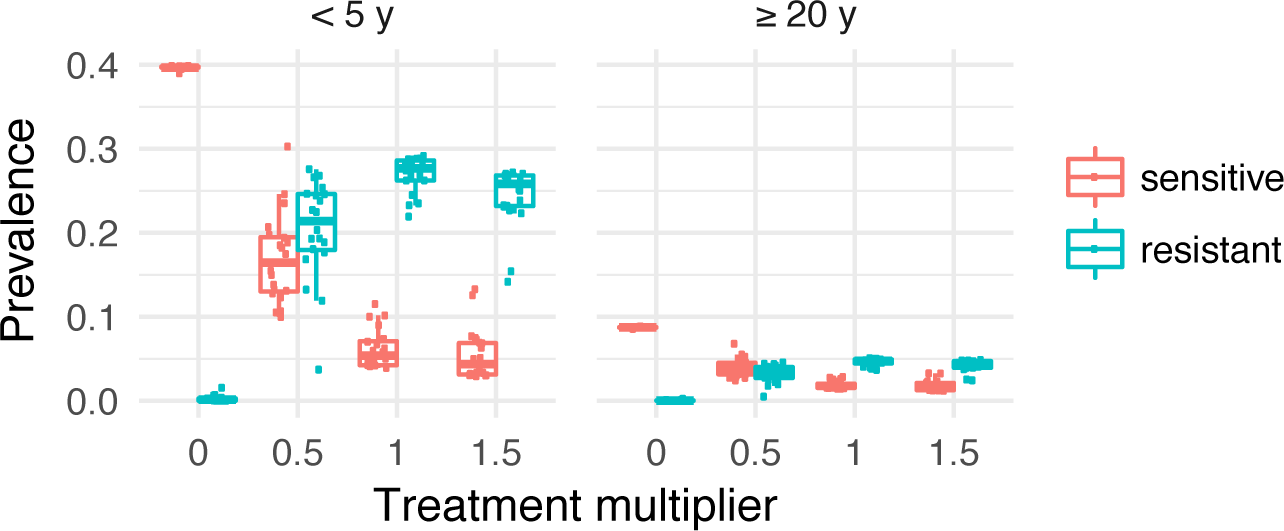
Prevalence of resistant and sensitive strains in young children (*<* 5 y) and adults (≥ 20y) in the model with age-assortative mixing (AAM), age-specific treatment (AST), and pseudo-spatial immigration (PSI). The ratio of the duration of resistant to sensitive strain’s durations in treated hosts (*γ*_r_) was set to 4, and the fitness cost of resistance was 4% (*ξ* = 0.96) and acted on duration. Each point shows the mean fraction of each serotype that is resistant from one replicate simulation. Means were obtained by averaging the last 50 years.

**Figure S3:**
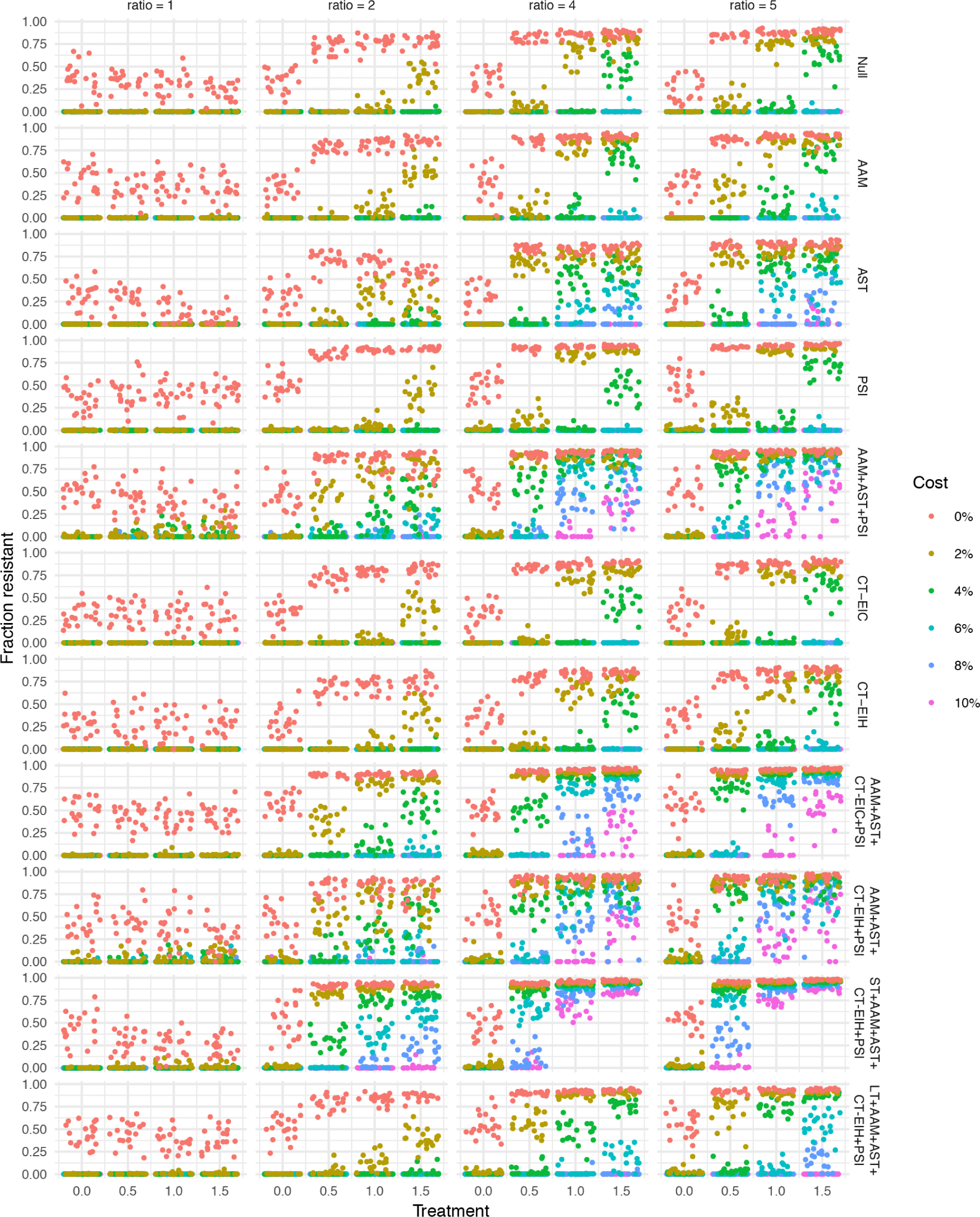
The fraction of resistant strains by treatment level and the ratio of resistant strains’ duration compared to sensitive strains’ in treated hosts. Each point corresponds to one simulation. AAM = age-assortative mixing, AST = age-specific treatment, PSI = pseudo-spatial immigration, CT = cotransmission, EIH = equally infectious hosts, EIC = equally infectious colonizations, trans = fitness cost in transmission instead of duration; ST = short treatment (high frequency), LT = long treatment (low frequency).

**Figure S4:**
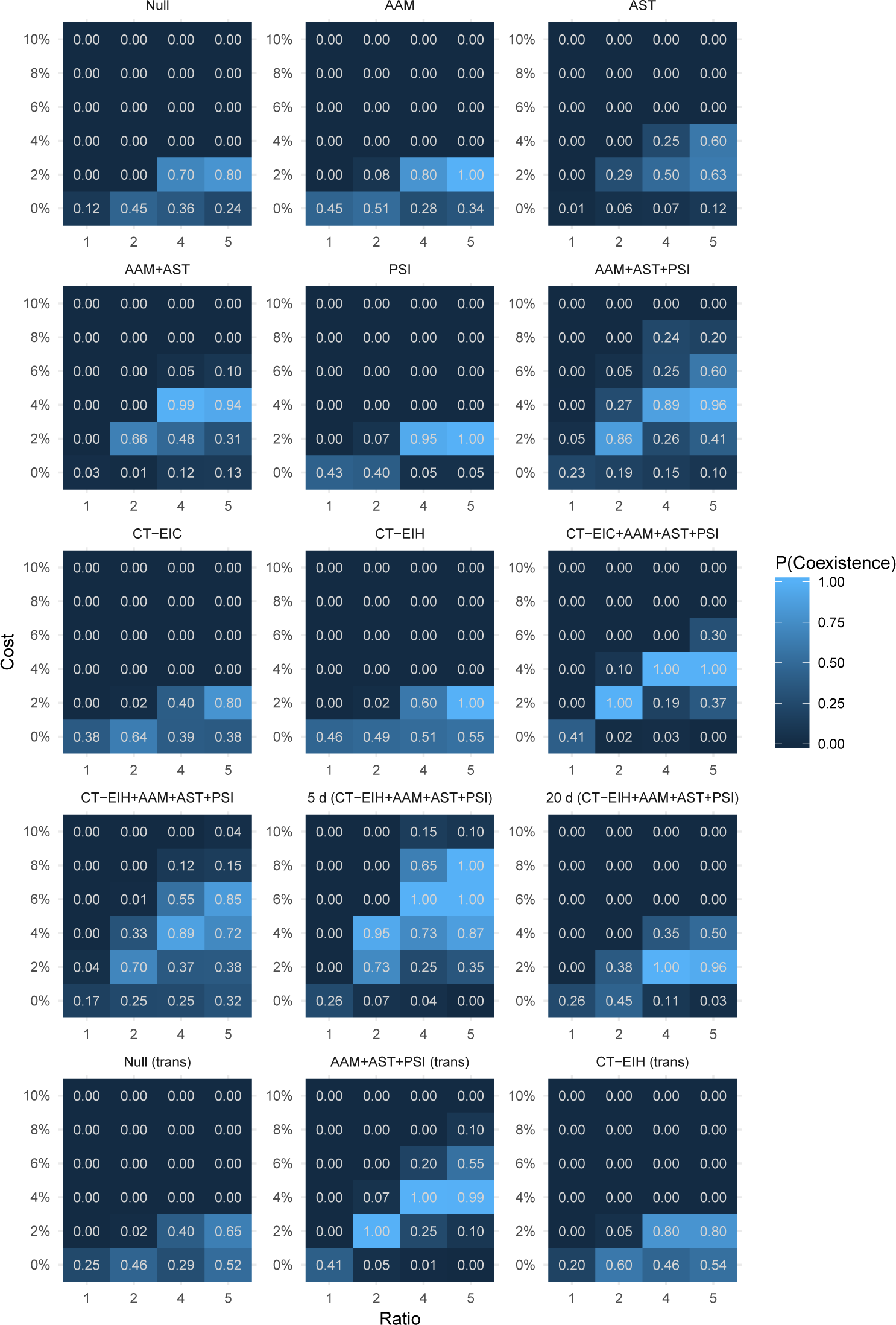
Fraction of replicate simulations producing coexistence for different fitness costs, ratios of the resistant to sensitive strain’s duration in treated hosts, and models. Coexistence is defined as a resistant fraction of 2%-98% and a ≥ 10% increase in the fraction resistant over 50% to 150% of typical treatment rates (see Fig. S3). AAM = age-assortative mixing, AST = age-specific treatment, PSI = pseudo-spatial immigration, CT = cotransmission, EIH = equally infectious hosts, EIC = equally infectious colonizations, trans = fitness cost in transmission instead of duration.

**Figure S5:**
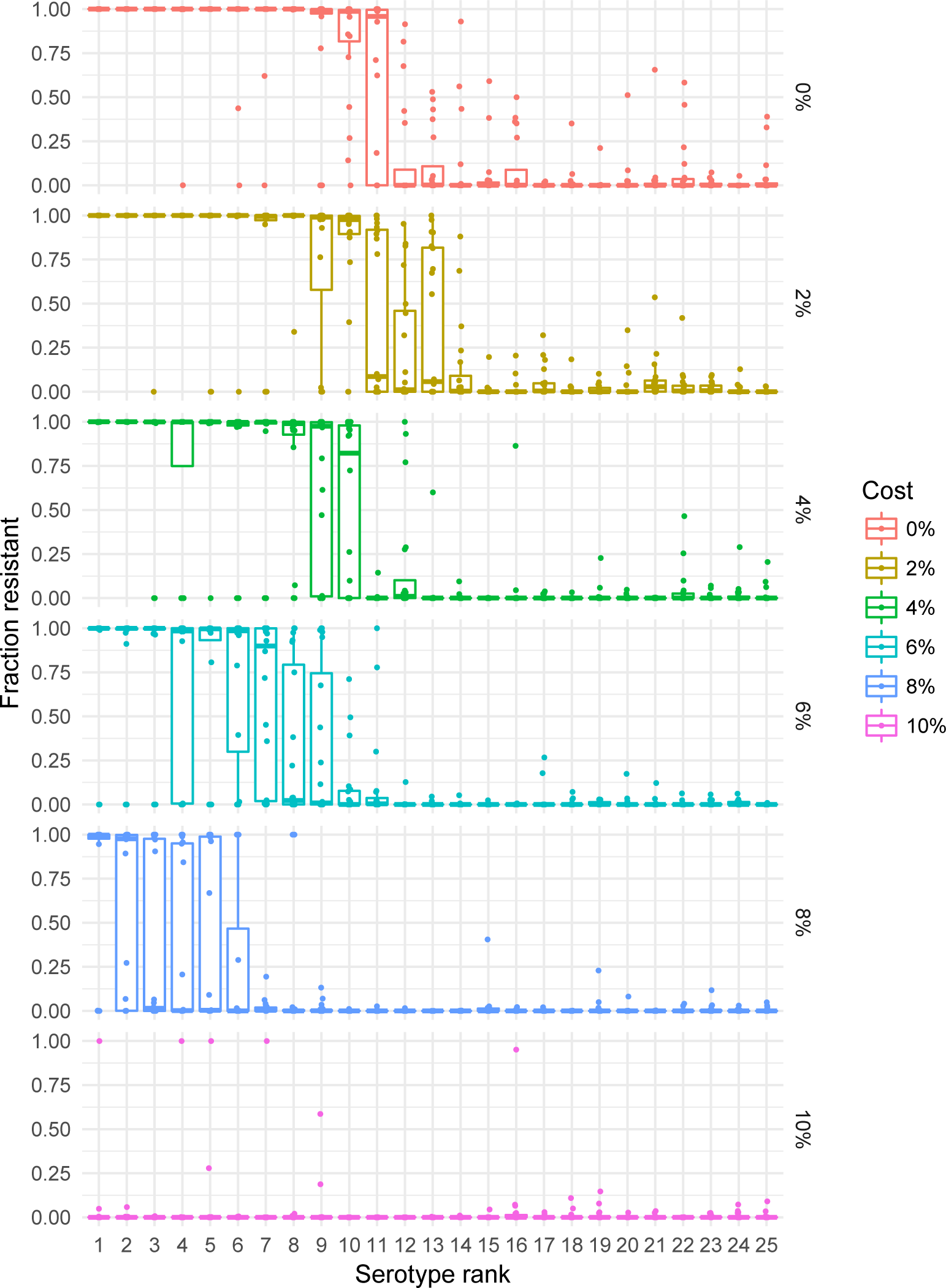
Resistance in adults (≥ 20 y) by serotypes’ fitness ranks for different costs of resistance in the model with age-assortative mixing (AAM), age-specific treatment (AST), pseudo-spatial immigration (PSI), and fitness cost in the duration of carriage. Treatment is held at 1.0, equivalent to U.S. rates. Each point shows the mean fraction of each serotype that is resistant from one replicate simulation. Means were obtained by averaging the last 50 years.

**Figure S6:**
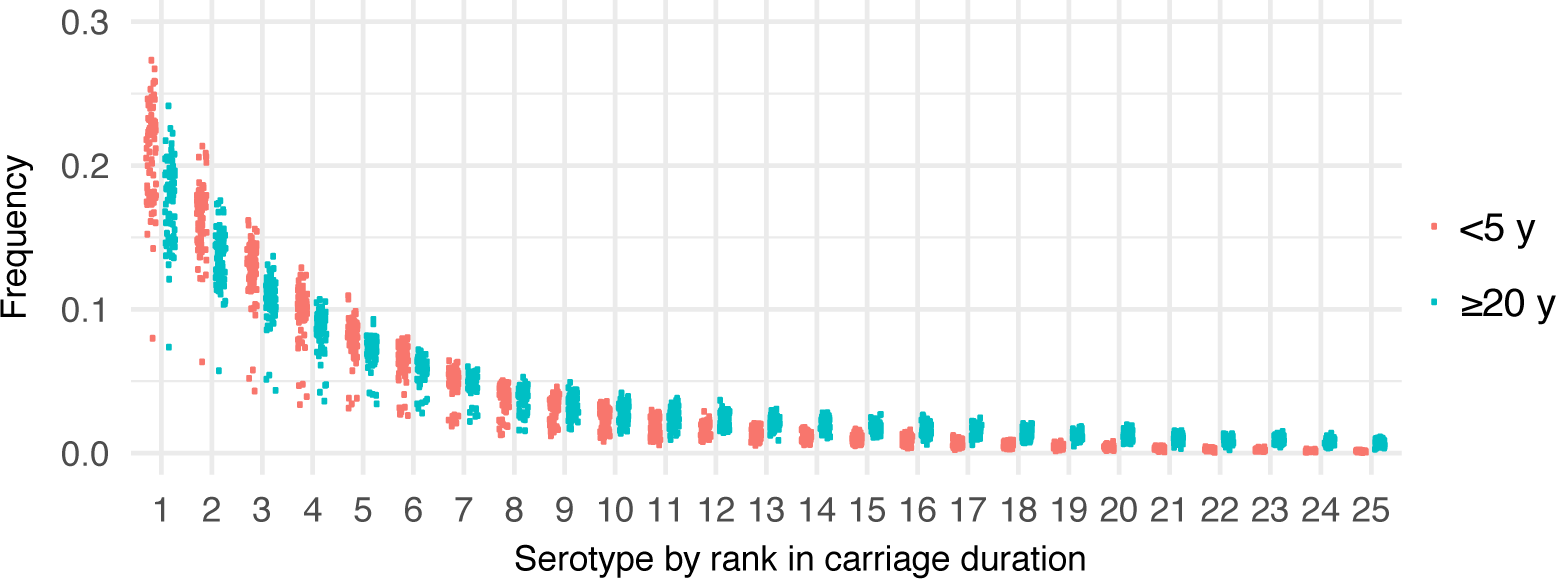
Frequencies of serotypes ranked by their carriage duration, from the longest-duration serotype (first) to the shortest-duration serotype (last), in children < 5 y and adults ≥ 20 y. Each point shows the mean frequency of each serotype (of all colonizations) from one replicate simulation of a model with age-assortative mixing (AAM) and age-structured treatment (AST) with treatment level 1 and a fitness cost of 2% (*ξ* = 0:98). Means were obtained by averaging frequencies from the last 50 years.

**Figure S7:**
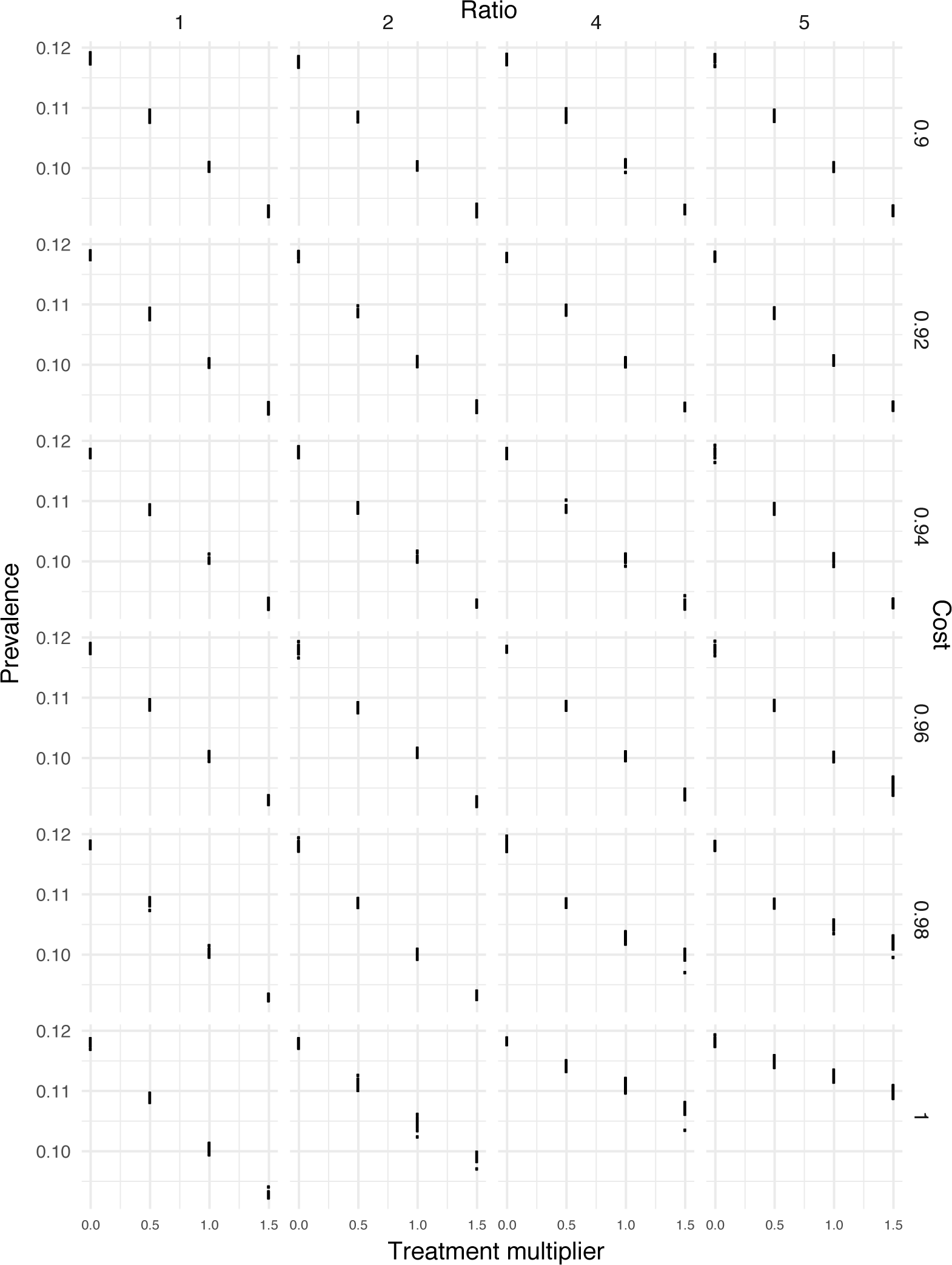
Mean carriage prevalence in the host population for the model with age-assortative mixing (AAM), age-specific treatment (AST), pseudo-spatial immigration (PSI), and the fitness cost of resistance in the duration of carriage. Each point shows the mean prevalence from one replicate simulation. Means were obtained by averaging the last 50 years.

**Figure S8:**
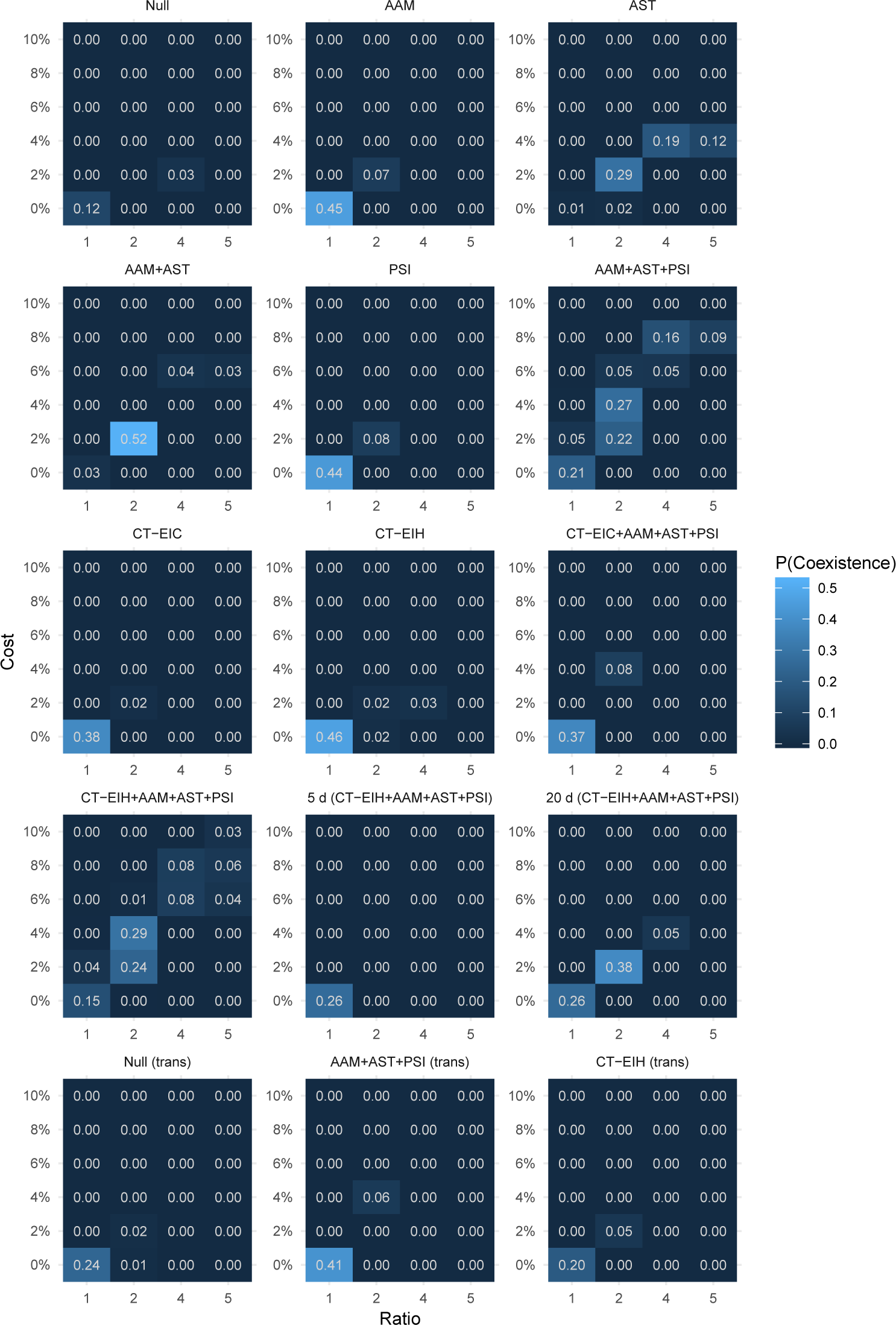
Fraction of replicate simulations producing coexistence for different fitness costs, ratios of the resistant to sensitive strain’s duration in treated hosts, and models. Coexistence is defined as a resistant fraction of 2%-60% and a ≥ 10% increase in the fraction resistant over 50% to 150% of typical treatment rates (see Fig. S3). AAM = age-assortative mixing, AST = age-specific treatment, PSI = pseudo-spatial immigration, CT = cotransmission, EIH = equally infectious hosts, EIC = equally infectious colonizations, trans = fitness cost in transmission instead of duration.

